# Hybrid assembly using ultra-long reads resolves repeats and completes the genome sequence of a laboratory strain of *Staphylococcus aureus* subsp. aureus RN4220

**DOI:** 10.1101/2021.06.25.449935

**Authors:** Suresh Panthee, Hiroshi Hamamoto, Atmika Paudel, Chikara Kaito, Yutaka Suzuki, Kazuhisa Sekimizu

## Abstract

*Staphylococcus aureus* RN4220 has been extensively used by staphylococcal researchers as an intermediate strain for genetic manipulation due to its ability to accept foreign DNA. Despite its wide use in laboratories, its complete genome is not available. In this study, we used the hybrid genome assembly approach using the minION long reads and Illumina short reads to sequence the complete genome of *S. aureus* RN4220. The comparative analysis of the annotated complete genome showed the presence of 39 genes fragmented in the previous assembly, many of which were located near the repeat regions. Using RNA-Seq reads, we showed that a higher number of reads could be mapped to the complete genome than the draft genome and the gene expression profile obtained using the complete genome also differs from that obtained from the draft genome. Furthermore, by comparative transcriptomic analysis, we showed the correlation between expression levels of staphyloxanthin biosynthetic genes and the production of yellow pigment. This study highlighted the importance of long reads in completing the microbial genomes, especially those possessing repetitive elements.

## Introduction

*Staphylococcus aureus* is a Gram-positive bacterium capable of opportunistic infections, which can sometimes be fatal. Genetic manipulation of *S. aureus* was limited until *S. aureus* strain RN4220 was obtained by chemical mutagenesis of *S. aureus* NCTC8325-4 strain[1]. NCTC8325-4 is a derivative of a clinical isolate NCTC8325 obtained by curing the three prophages Φ11, Φ12, and Φ13[2]. Therefore, both NCTC8325-4 and RN4220 lack the three prophages. In addition, RN4220 can accept foreign DNA and is characterized by a mutation in the *sauI hsdR* gene belonging to the restriction-modification system[3]. Due to this property, RN4220 is routinely used in the laboratories as an intermediate for genetic manipulation; plasmids from *Escherichia coli* are electro-transformed into RN4220, and the plasmids from RN4220 are then transformed to another *S. aureus* strain by suitable methods such as phage transduction.

Despite its wide use, the complete genome of this strain is not available. With the recent development in next-generation sequencing technologies, there have been attempts to sequence the genome. Apart from our assembly, there are two deposited assemblies of RN4220 in NCBI. The first assembly was done in 2011 using Illumina GA II[4]( accession: GCA_000212435.2), and the second was performed in 2020 using BGISeq (accession: GCA_011751615.1), which generated 118 and 27 contigs, respectively. Whereas the assemblies primarily provided valuable information regarding the genetic make-up of this strain, we still need a complete genome sequence to make the most out of this laboratory strain. Short reads sequencing of the genome can be attributed to the large number of contigs generated from these assemblies. Short read assemblies are challenged by the presence of identical sequences at more than one locus of the chromosome called the repetitive DNA sequences, or repetitive elements or repeated regions. Larger organisms such as eukaryotes have many repetitive elements throughout the genome; for instance, nearly half of the human genome consists of repetitive elements[5]. Bacteria such as *Orientia tsutsugamushi* possess 37% repetitive elements throughout its genome[6]. In this study, by hybrid assembly using both long and short reads, we completed the genome of RN4220. Upon further analysis, we found many repetitive elements and several fragmented genes in the previous assembly. Our approach using long reads was suitable for covering those repetitive regions, and we were successful in obtaining a complete genome.

## Materials and Methods

### Genome sequencing, assembly, and annotation

*S. aureus* RN4220 was routinely cultured on tryptic soy broth at 37°C without antibiotic selection. Genome sequencing and assembly was performed as previously explained[7–10]. Briefly, genomic DNA was isolated from overnight culture using Qiagen DNA-blood Mini Kit (Qiagen, Hilden, Germany) and lysostaphin for bacterial lysis. Construction of short-read single-end libraries and sequencing was performed using Illumina HiSeq2000 (Illumina, San Diego, CA, USA)[11]. MinION long reads sequencing was performed using 1 μg genomic DNA. Hybrid error correction of the long reads was performed by LoRDEC[12] using the short reads, and final assembly of the circular chromosome was performed using Flye 2.3.3[13]. Short reads were then mapped to the chromosome, and the consensus was generated to obtain the final assembly. The final assembly was then annotated using the NCBI Prokaryotic Genome Annotation Pipeline (PGAP).

### Comparative genomic analysis

The complete genome sequence of the parent strain NCTC8325 and two draft assemblies of RN4220 strain were downloaded from NCBI. The draft assemblies were first aligned using Mauve Contig Mover[14]. The ordered contigs were then submitted to the CLC Genomics workbench for whole-genome alignment. To identify the genes fragmented in the previous assembly, the annotations were checked manually. Repeat finding of the genome was performed using Unipro UGENE v.39.0[15].

### Staphyloxanthin biosynthesis pathway analysis

Raw RNA-Seq reads for *S. aureus* RN4220, NCTC8325, and SH1000 were downloaded from NCBI SRA. The SRA accession numbers and BioProject details are summarized in **Table 1**. The reads were then mapped to their respective complete genome using the CLC Genomic Workbench ver 20.0.4 (CLC bio, Aarhus, Denmark). Since the complete genome sequence of the SH1000 strain is not publicly available, the SH1000 reads were mapped to the NCTC8325 genome. Transcripts Per Million (TPM), a sequence depth normalized indicator for expression analysis, was used to compare the expression level among the study samples.

**Table 1:**
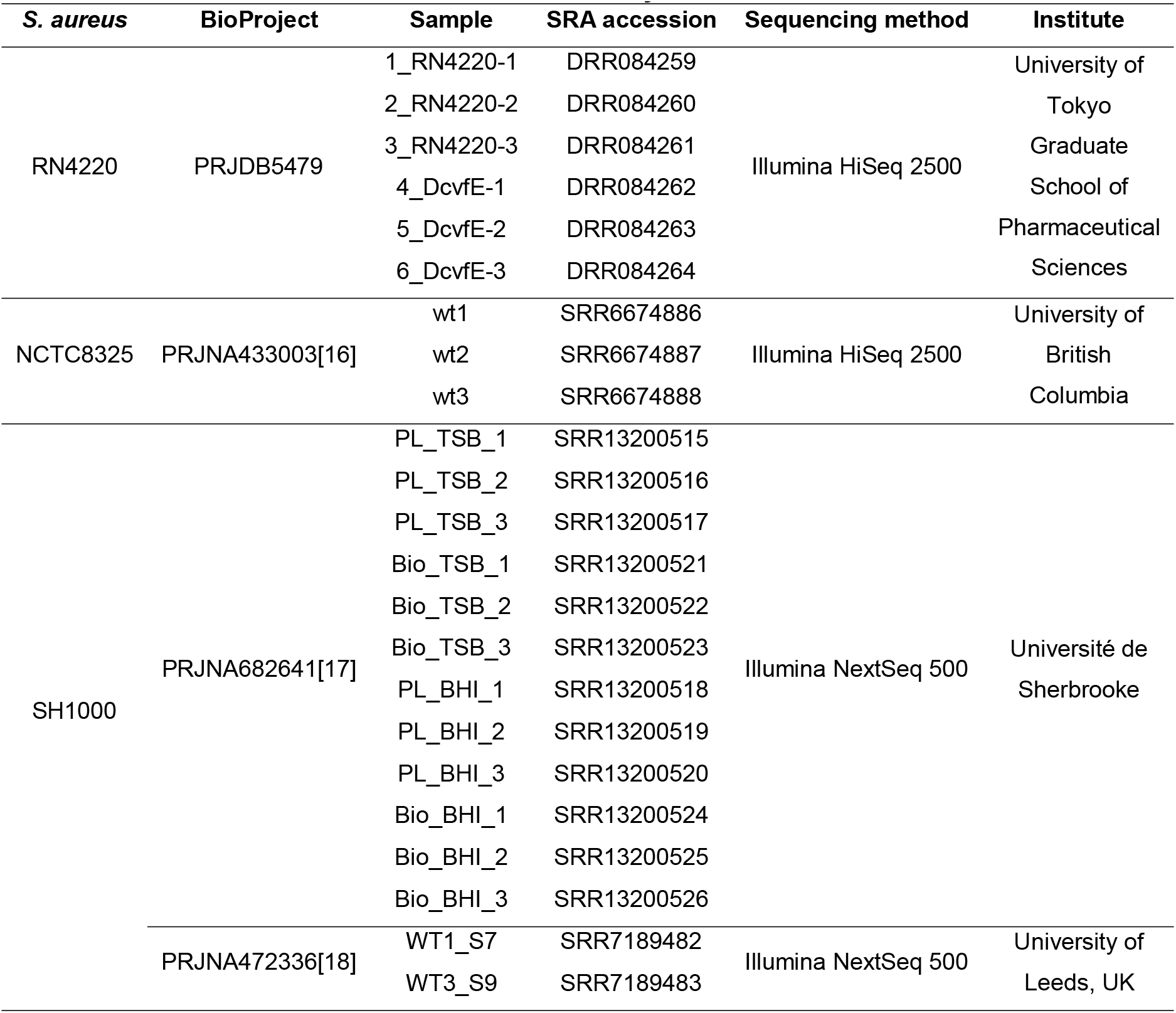
SRA accession numbers used in this study.

| *S. aureus* | BioProject | Sample | SRA accession | Sequencing method | Institute |
| --- | --- | --- | --- | --- | --- |
| RN4220 | PRJDB5479 | 1_RN4220-1 | DRR084259 | Illumina HiSeq 2500 | University of<br>Tokyo<br>Graduate<br>School of<br>Pharmaceutical<br>Sciences |
|  |  | 2_RN4220-2 | DRR084260 |  |  |
|  |  | 3_RN4220-3 | DRR084261 |  |  |
|  |  | 4_DcvfE-1 | DRR084262 |  |  |
|  |  | 5_DcvfE-2 | DRR084263 |  |  |
|  |  | 6_DcvfE-3 | DRR084264 |  |  |
| NCTC8325 | PRJNA433003[16] | wt1 | SRR6674886 | Illumina HiSeq 2500 | University of<br>British<br>Columbia |
|  |  | wt2 | SRR6674887 |  |  |
|  |  | wt3 | SRR6674888 |  |  |
| SH1000 | PRJNA682641[17] | PL_TSB_1 | SRR13200515 | Illumina NextSeq 500 | Université de<br>Sherbrooke |
|  |  | PL_TSB_2 | SRR13200516 |  |  |
|  |  | PL_TSB_3 | SRR13200517 |  |  |
|  |  | Bio_TSB_1 | SRR13200521 |  |  |
|  |  | Bio_TSB_2 | SRR13200522 |  |  |
|  |  | Bio_TSB_3 | SRR13200523 |  |  |
|  |  | PL_BHI_1 | SRR13200518 |  |  |
|  |  | PL_BHI_2 | SRR13200519 |  |  |
|  |  | PL_BHI_3 | SRR13200520 |  |  |
|  |  | Bio_BHI_1 | SRR13200524 |  |  |
|  |  | Bio_BHI_2 | SRR13200525 |  |  |
|  |  | Bio_BHI_3 | SRR13200526 |  |  |
|  | PRJNA472336[18] | WT1_S7 | SRR7189482 | Illumina NextSeq 500 | University of<br>Leeds, UK |
|  |  | WT3_S9 | SRR7189483 |  |  |

## Results and Discussions

### Completion of the genome sequence of *S. aureus* RN4220

Previous attempts to sequence the RN4220 genome used the sequencers that produced short reads. *S. aureus* NCTC8325, the parent strain of RN4220, also possesses repetitive elements, which create a challenge in assembling the genome using short reads. To cover the repetitive elements while sequencing a genome, either the reads longer than the repetitive elements or alternative approaches to overcoming this problem are essential. In this study, we took the advantage of our hybrid genome assembly approach[7–10] using long reads from ONT MinION and short reads from Illumina to complete the RN4220 genome. As high-quality long reads are much crucial than coverage, we could complete the genome of this bacterium with low coverage long reads, and high coverage short reads. The summary of reads obtained from both the MinION and Illumina platforms is shown in **Figure 1A, B**.

**Figure 1.**
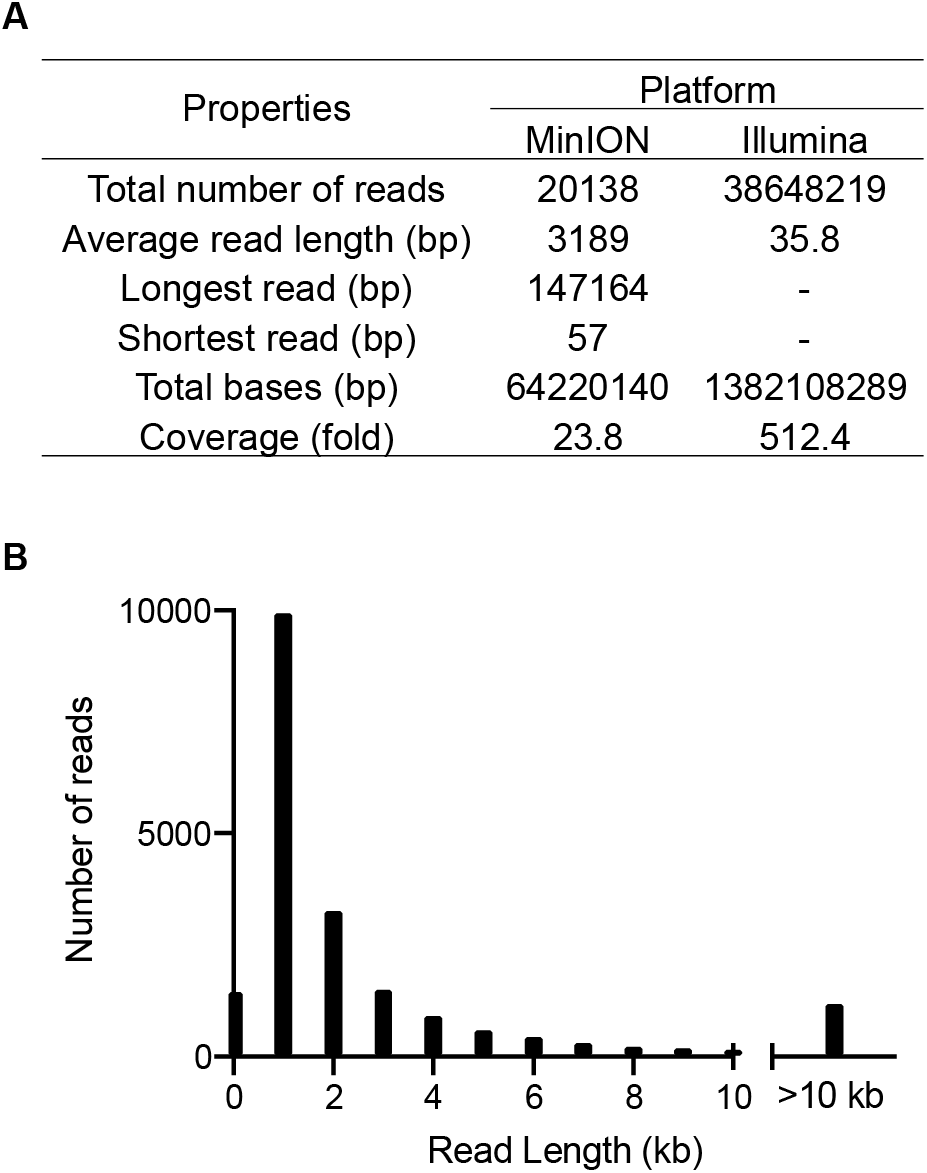
Genome sequencing of *S. aureus* RN4220 using hybrid-genome assembly approach. (**A**) Summary of sequence reads from MinION and Illumina sequencers. (**B**) Histogram of read length obtained from the MinION sequencer.

The complete genome sequence of *S. aureus* RN4220 was 2.7 Mb in length, harbored 2654 genes, including 19 rRNAs and 59 tRNAs (**Table 2**). To identify the genomic difference among the parent strain and RN4220 assemblies, we performed the whole genome alignment of the parent strain NCTC8325[19] (assembly accession: GCA_000013425.1). We found that large regions from NCTC8325 were deleted in the RN4220 genome, which has also been noted by the previous analysis using the draft genome [4].

**Table 2:** Analysis and comparison of general features of the current, complete *S. aureus* RN4220 genome with previous draft assemblies.

| Features | Current assembly<br>GCA_018732165.1 | GCA_011751615.1 | GCA_000212435.2[4] |
| --- | --- | --- | --- |
| Assembly release date | 2021/06/07 | 2020/03/25 | 2011/05/05 |
| Sequencing technology | ONT minION<br>Illumina HiSeq | BGIseq | Illumina GA II |
| Genome coverage | ONT minION: 23x<br>Illumina HiSeq: 512x | 372x | 77x |
| Total Sequence length (bp) | 2,697,195 | 2,657,542 | 2,663,395 |
| No of contigs | 1 | 27 | 118 |
| Contig N50 | - | 174,720 | 80,460 |
| Contig L50 | - | 4 | 13 |
| Protein coding genes | 2508 | 2517 | 2540 |
| rRNA genes | 78 | - | - |
| CDS | 2572 | 2571 | 2604 |
| Gene | 2654 | 2626 | 2661 |
| Misc. binding | 3 | 3 | 3 |
| Misc. feature | 3 | 3 | 3 |
| ncRNA | 3 | 3 | 3 |
| Regulatory | 10 | 10 | 10 |
| rRNA | 19 | - | 8 |
| tmRNA | 1 | 1 | 1 |
| tRNA | 59 | 51 | 45 |

**Figure 2.**
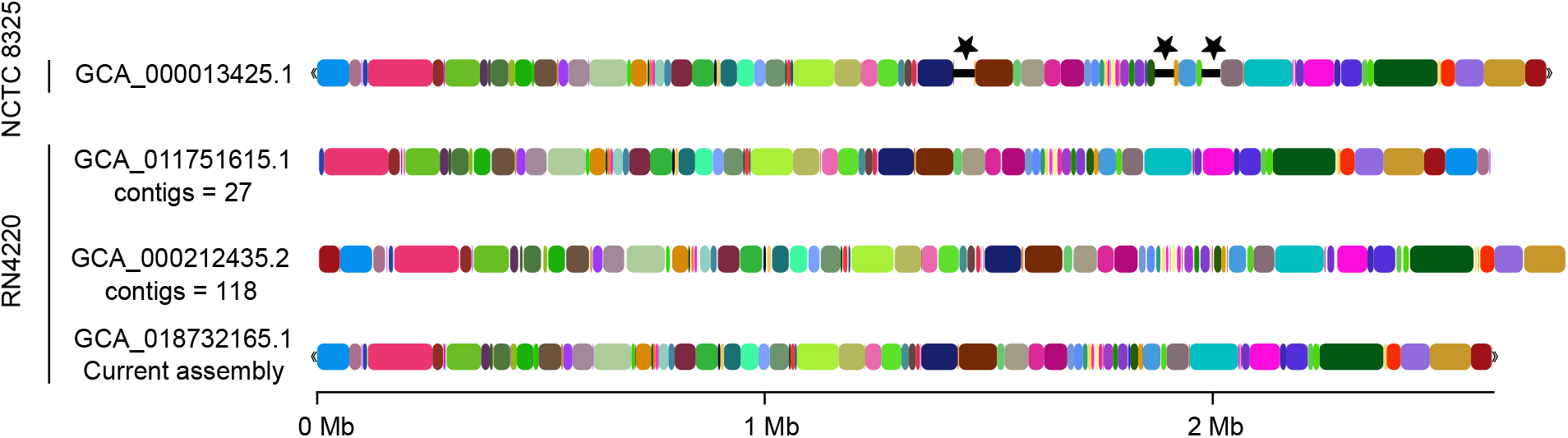
Comparison of *S. aureus* RN4220 genome. Whole-genome alignment of the parent strain NCTC8325 with RN4220 assemblies. The three regions indicated by the stars were present in the NCTC8325 but not in the RN4220 genomes. The homologous regions were randomly colored for ease of distinction. The assembly GCA_000212435.2 appears longer due to the large number of contigs.

### Identification of fragmented genes and presence of repetitive elements

We compared our current assembly with the first genome assembly and found that many genes were fragmented and possibly not detected earlier. We found 39 new genes among the fragmented regions. Some notable genes identified included KMZ21_00310: *spa*; KMZ21_02625: *sdrC*; KMZ21_02630: *sdrD*; KMZ21_06705: *ebh*; and KMZ21_08630: *splF.* As these proteins are known to be involved in *S. aureus* pathogenesis through processes such as immune evasion, adhesion, complement resistance, and substrate acquisition, it is speculated that the virulence potential, based on the draft genome sequence, might have been overlooked. Furthermore, we found that 14 of these genes were located around the repetitive elements, suggesting the importance of long reads in the genome assembly (**Table 3**).

**Table 3:**
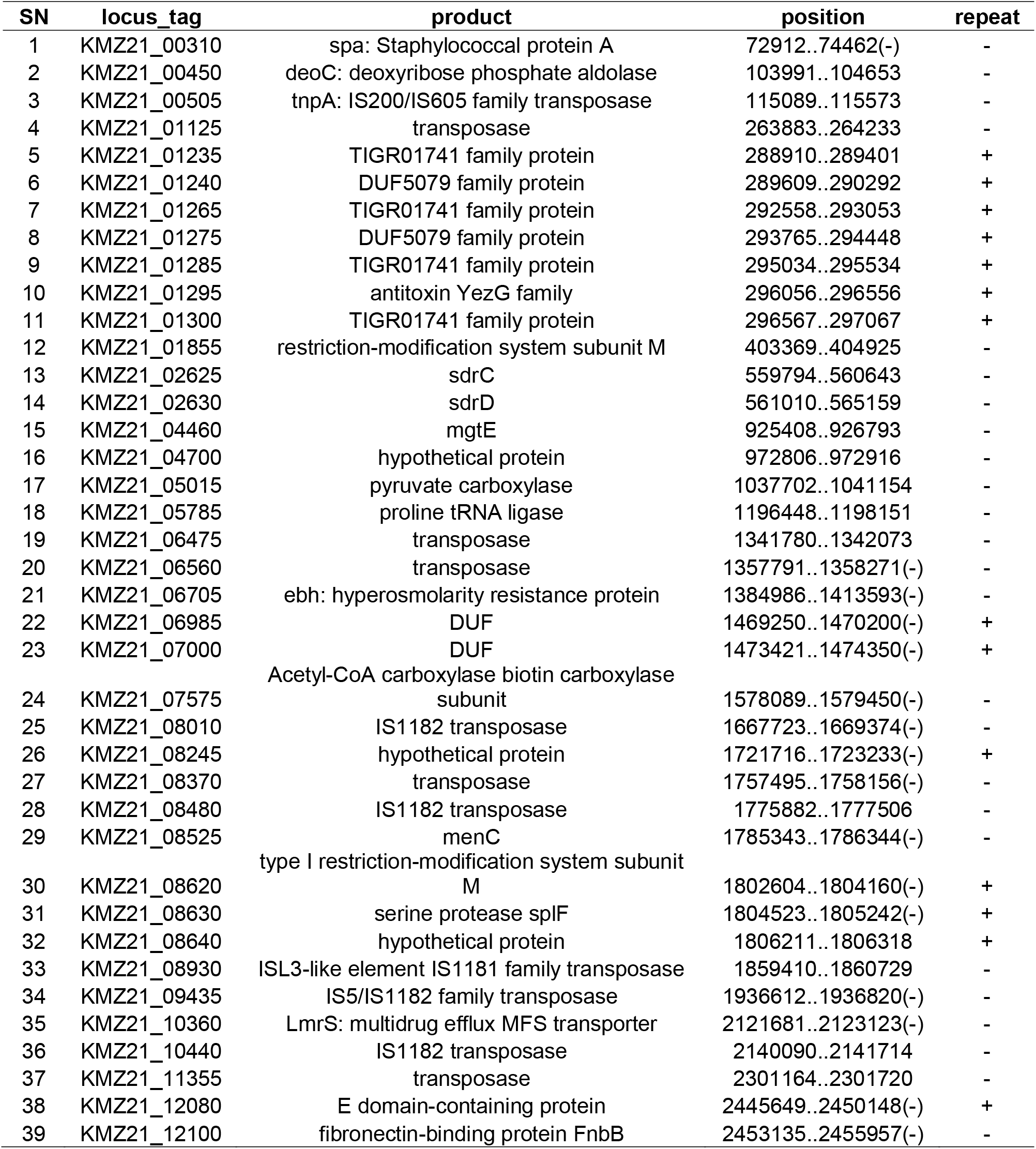
Fragmented genes in the previous assembly [4], identified through the complete genome analysis. (-) in the position, column indicates gene in complementary strand, and + or - sign in the repeat column indicates a presence or absence of repetitive elements in the proximity of the gene, respectively.

### Complete genome facilitates the RNA-Seq analysis

Using the publicly available data, we calculated the mapping of short RNA-Seq reads and fold expression of genes and compared the data with the complete and draft RN4220 genomes. We found that the number of reads mapped to the complete genome was slightly higher in the complete genome than that of the draft genome (**Figure 3A**). It is well known that the scaffolds in most draft genomes contain gaps[20] which may sometimes lead to the difference in the mapping of the reads and might be misleading while interpreting results. Therefore, we can expect that the increased mapping could be due to the mapping of the additional reads in the “gap” region that lied in between the contigs in the draft genome. Interestingly, the number of reads mapped specifically to the genome drastically reduced in the complete genome (**Figure 3A**). This could be because of the resolved repeat and duplicate regions in the complete genome, which appeared as a single contig in the draft genome. We found more than 80% of the reads were matched to six positions in the complete genome, which in the case of the draft genome was one (**Figure 3B**). To compare the difference between the gene expression analysis obtained using a complete genome and draft genome, we used RNA-seq reads from RN4220 wild-type and ΔKMZ21_09125 (SA1684), involved in virulence of *S. aureus* strains[21]. Using a volcano plot (**Figure 3C**), we could see a vast difference in the differential expression of the genes when using the complete and the draft genome. Besides, the lack of the RN4220 complete genome required researchers to analyze RNA-Seq reads by mapping to the NCTC8325 genome[21, 22], where the results should be carefully interpreted, considering the differences among NCTC8325 and RN4220 genome sequences. In summary, these suggested the importance of a complete genome for omics-based analysis.

**Figure 3:**
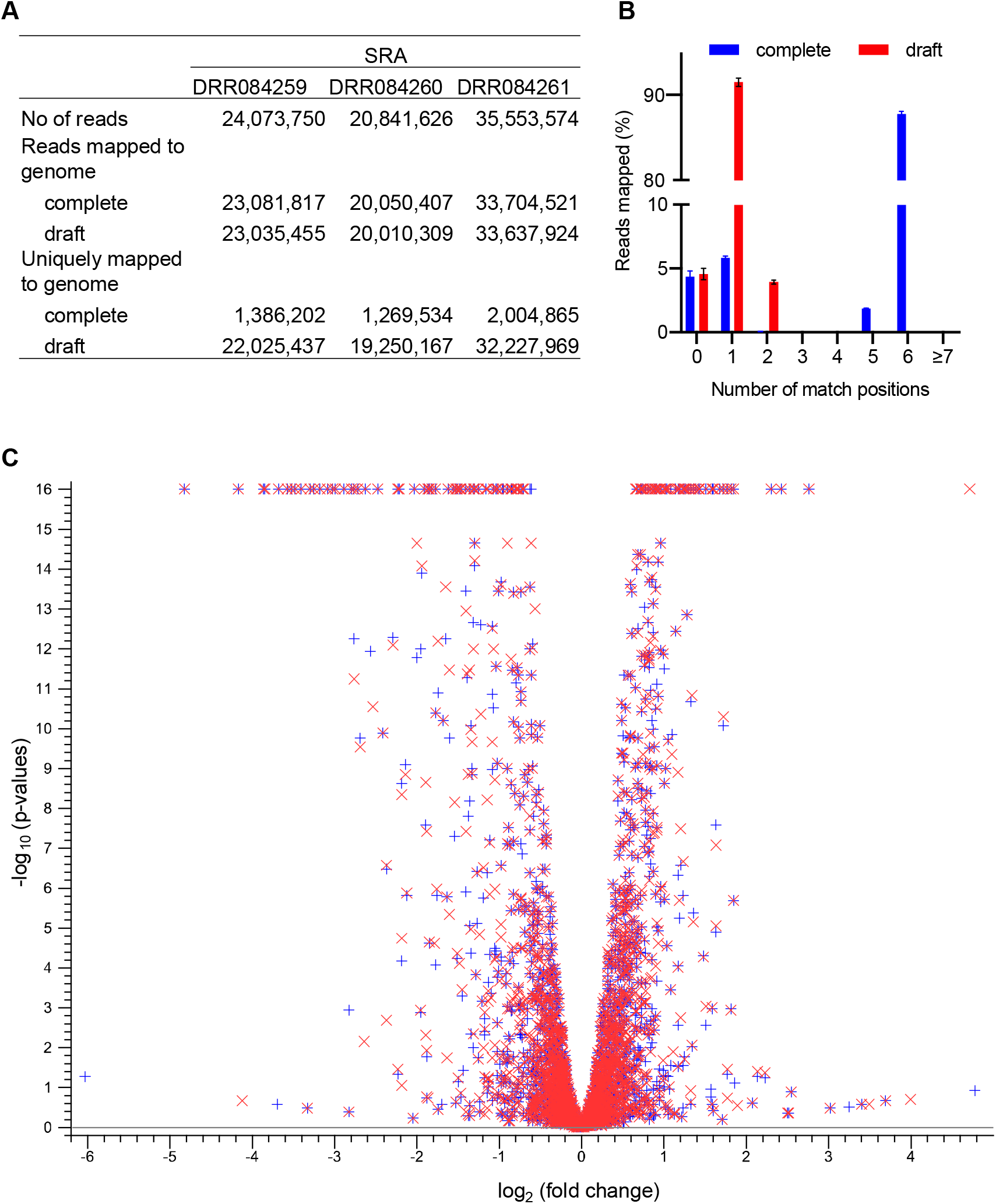
Comparison of complete (current) and draft[4] genome assemblies to analyze RNA-Seq results. Mapping (A) and match specificities (B) of RN4220 wild-type reads to the genome. (C) The differential expression of the genes between Δ*cvfE* and wild-type strains using complete and draft genome (+: complete genome; x: draft genome).

### Downregulation of RN4220 genes involved in staphyloxanthin biosynthesis

*S. aureus* strains are usually distinguishable from other bacteria due to their yellow color, which is because of the production of yellow pigment staphyloxanthin. However, the RN4220 strain does not give the yellow pigmentation, hence produces very little or no staphyloxanthin. Staphyloxanthin biosynthetic genes are located in an operon *crtMNOPQ* (**Figure 4A**)[23, 24] which is dependent upon the sigma factor B (SigB)[25]. SigB falls in an operon *rsbUVWSigB,* where RsbU and RsbV are the activators, and RsbW is the repressor of SigB[26–28]. It has been known that SigB is also controlled by YjbH[29] and CspA[30]. The parent strain of RN4220, NCTC8325, has a reduced ability to produce staphyloxanthin[31] and this is attributed to a deletion of 11 bp in the *rsbU* gene. Since staphyloxanthin production is more pronouncedly decreased in RN4220 compared with NCTC8325 and NCTC8325-4[31], we expected that the RN4220 strain might have some further alterations within these two operons. We aligned the amino acid sequences of the genes and found that these operons were conserved. Next, we aimed to examine the difference at the gene expression level. We looked for the raw RNA-Seq reads in NCBI SRA for NCTC8325, RN4220, and SH1000 strains. The *S. aureus* SH1000[32] strain is a *rsbU*^+^ derivative of NCTC8325 and has the ability to produce staphyloxanthin. The reads were then mapped to complete genomes, and expression was analyzed using transcripts per million (TPM). We found that the *crt* operon was dramatically downregulated in RN4220, consistent with its pigment-less phenotype (**Figure 4B**). As we did not find a genetic level changes within the *crt* operon, it is expected that some other regulatory factors might play a role in the observed difference in staphyloxanthin production.

**Figure 4:**
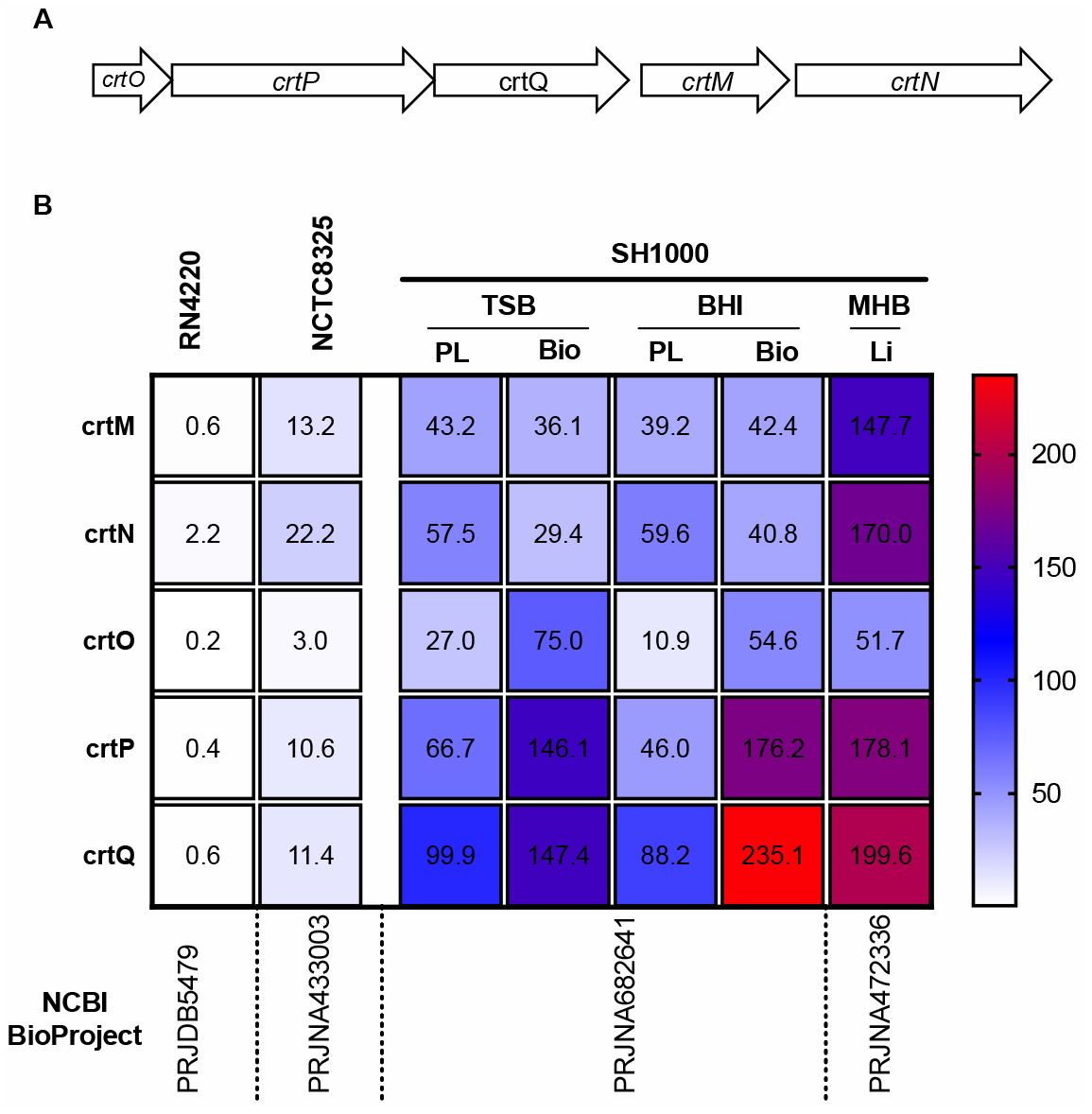
Staphyloxanthin biosynthetic gene cluster (A) and expression of the genes among NCTC8325, and SH1000, the *rsbU* repaired NCTC8325-4 strain (B). The short reads from the NCBI database (indicated Bioproject) were downloaded, mapped to respective genomes, and expression values, expressed as TPMs, were calculated using CLC Genomics Workbench. The mean of TPM values is shown. For RN4220, wild-type data was used. PL: Planktonic growth; Bio: Biofilm growth; Li: liquid culture.

## Conclusion

In this study, we completed the genome of a popular laboratory strain RN4220 for the first time. We provided an example of the importance of long reads in completing genomes containing repetitive elements, which is not possible by usual short-read sequences. The availability of the complete genome of this widely used strain is expected to serve as a platform for further genetic manipulation in a defined manner and robust omics-based analysis. In addition, we found that although the staphyloxanthin gene cluster was intact in RN4220, transcription of the operon was weak, resulting in a dramatic decrease in staphyloxanthin production and, hence, its pigment-less phenotype. Overall, the findings of this study provide valuable information on *S. aureus* RN4220 strain by completing its genome, which will help interpret results in a defined manner by reducing biases and broadening our understanding of the genetic basis of various phenotypes.

## Acknowledgments

This work was supported by JSPS KAKENHI Grant Numbers 19K16653JP to A.P., 19K07140JP to H.H., 21H02733JP to K.S., and 15H05783 to K.S. and H.H., and TBRF, IFO fellowships to S.P.

## Data availability

The complete genome of *S. aureus* RN4220 has been deposited to NCBI GenBank with accession CP076105.

## Author contributions

S.P. and H.H. conceived the idea. S.P. and A.P. performed long-read sequencing, assembled, analyzed the genome, and wrote the manuscript. Y.S. and H.H. performed short-read sequencing. C.K. performed RNA-seq analysis of RN4220 wild-type and ΔKMZ21_09125. H.H., C.K. and K.S. provided critical discussion for writing the manuscript. All authors contributed critically to the interpretation of the results and gave final approval for publication.

